# Genetic diversity of Collaborative Cross mice controls viral replication, clinical severity and brain pathology induced by Zika virus infection, independently of *Oas1b*

**DOI:** 10.1101/677484

**Authors:** Caroline Manet, Etienne Simon-Lorière, Grégory Jouvion, David Hardy, Matthieu Prot, Marie Flamand, Jean-Jacques Panthier, Anavaj Sakuntabhai, Xavier Montagutelli

## Abstract

The explosive spread of Zika virus (ZIKV) has been associated with major variations in severe disease and congenital afflictions among infected populations, suggesting an influence of host genes. We investigated how genome-wide variants could impact susceptibility to ZIKV infection in mice. We first describe that the susceptibility of *Ifnar1* knockout mice is largely influenced by their genetic background. We then show that the broad genetic diversity of Collaborative Cross mice, which receptor to type I interferon (IFNAR) was blocked by anti-IFNAR antibody, expressed phenotypes ranging from complete resistance to severe symptoms and death with large variations in the peak and rate of decrease of plasma viral load, in brain viral load, in brain histopathology and in viral replication rate in infected cells. Differences of susceptibility between CC strains were correlated between Zika, Dengue and West Nile viruses. We identified highly susceptible and resistant mouse strains as new models to investigate the mechanisms of human ZIKV disease and other flavivirus infections. Genetic analyses revealed that phenotypic variations are driven by multiple genes with small effects, reflecting the complexity of ZIKV disease susceptibility in human population. Notably, our results rule out a role of the *Oas1b* gene in the susceptibility to ZIKV. Altogether, this study emphasizes the role of host genes in the pathogeny of ZIKV infection and lays the foundation for further genetic and mechanistic studies.

**IMPORTANCE:** In recent outbreaks, ZIKV has infected millions of people and induced rare but potentially severe complications, including Guillain-Barré syndrome and encephalitis in adults. While several viral sequence variants were proposed to enhance the pathogenicity of ZIKV, the influence of host genetic variants in the clinical heterogeneity remains mostly unexplored. We have addressed this question using a mouse panel which models the genetic diversity of human population and a ZIKV strain from a recent clinical isolate. Through a combination of *in vitro* and *in vivo* approaches, we demonstrate that multiple host genetic variants determine viral replication in infected cells, and clinical severity, kinetics of blood viral load and brain pathology in mice. We describe new mouse models expressing high susceptibility or resistance to ZIKV and to other flaviviruses. These models will facilitate the identification and mechanistic characterization of host genes that influence ZIKV pathogenesis.

---

Zika virus (ZIKV) is a mosquito-borne flavivirus isolated in 1947 from a febrile rhesus monkey in Uganda (1). Until 2007, ZIKV had circulated in Africa and Asia causing mild flu-like syndromes, with rare reported clinical cases (2). However, during recent epidemics, ZIKV infection triggered severe complications including Guillain-Barré syndrome and encephalitis in adults (3, 4), and congenital malformations in fetuses of infected pregnant women (5, 6). Viral mutations may have contributed to ZIKV enhanced pathogenicity (7, 8) but only partly explain the variable proportions of symptomatic infections (9) and the increased incidence of congenital Zika syndrome (CZS) in Polynesia (10) and Brazil (11), suggesting a role for host genetic variants. Recent evidence indicates that the regulation of innate immunity genes is driven by host genetic background in human fetal brain-derived neural stem cells (hNSCs) infected *in vitro* with ZIKV (12). Additionally, the analysis of pairs of dizygotic twins exposed to ZIKV during pregnancy and discordant for CZS suggests multigenic host susceptibility to ZIKV-induced brain malformations (13).

Multiple mouse models have been proposed to decipher the mechanisms of ZIKV disease pathogenesis (14, 15). These models allow the investigation of several key features of human infection, such as neuronal damage (16, 17), sexual and vertical transmission (18–21), fetal demise and CZS (22–25). However, while non-structural ZIKV proteins efficiently inhibit innate antiviral responses in humans (26, 27) allowing viral replication, ZIKV replicates poorly in wild-type mice due to its NS5 protein’s inability to antagonize STAT2 and type I interferon (IFN) response as in humans (28). Effective systemic infection in mice occurs when this response is abrogated by genetic inactivation of the *Ifnar1* gene (29) or by blocking the type I IFN receptor with the MAR1-5A3 monoclonal antibody (mAb) (30, 31). So far, host genetic factors involved in mouse susceptibility to ZIKV infection have been investigated mainly through reverse genetic approaches, by studying the consequences of genetic ablation of specific genes, such as innate or adaptive immunity genes (29, 32–35). While these models have contributed to our understanding of the mechanisms of ZIKV disease, they do not model the simultaneous contribution of variants in multiple pathways, as would most likely be observed in the natural population. A recent study has reported strain-specific differences in the susceptibility to neonatal ZIKV infection across four mouse laboratory strains, affecting neuropathology and behavior in adulthood (36). More extensive studies investigating the role of genome-wide genetic variations on the susceptibility to ZIKV infection are needed, using mouse models that accurately reflect the phenotypic and genetic diversity of the human population (37).

In this study, we addressed this question using the two types of susceptible mouse models. First, since the phenotype of a single gene modification often varies under the influence of modifier genes (38, 39), we assessed the effect of host genetic background on the susceptibility of *Ifnar1*-deficient mice. We then investigated the impact of host genetic diversity on the susceptibility to ZIKV infection in the Collaborative Cross (CC), a panel of recombinant inbred mice produced by a systematic cross between eight founder inbred strains, including five classic laboratory strains and three wild-derived strains (40). The founder strains capture approximately 90% of the genetic variants present in the *Mus musculus* species (41) and the resulting CC strains, which segregate an estimated 45 million polymorphisms, have more genetic diversity than the human population (42). Extensive variations in pathogenic phenotypes have been previously reported in the CC panel after viral (43–50), bacterial (51, 52) and fungal (53) infections, demonstrating that this resource is ideally suited for investigating the role of host genetic variants in the pathophysiology of infectious diseases (54).

Susceptibility to ZIKV in *Ifnar1*-deficient mice was strongly influenced by the genetic background, with practical implications for virology studies and allowing for future identification of modifier genes. The challenge of 35 immunocompetent CC strains with ZIKV after MAR1-5A3 mAb treatment allowed efficient viral replication. We show that genetic diversity in the CC panel enabled large variations in the clinical severity of ZIKV disease, in the peak and kinetics of plasma viral load and in the severity of ZIKV-induced brain pathology. Genetic diversity also resulted in differences in the permissiveness of CC mouse cells to viral replication which likely contributes to the *in vivo* phenotypic range. In a subset of CC strains, we found correlated differences of susceptibility to ZIKV, dengue virus (DENV) and West Nile virus (WNV), suggesting shared underlying mechanisms. We identified highly susceptible and resistant mouse strains as new models to investigate the mechanisms of human ZIKV disease and other flavivirus infections. Finally, genetic analysis revealed that susceptibility to ZIKV in the CC is driven by multiple loci with small individual effects, and that *Oas1b*, a major determinant of mouse susceptibility to WNV, is not involved.

## RESULTS

### Genetic background controls the susceptibility of *Ifnar1*-deficient mice to ZIKV

Many studies have used *Ifnar1* knock-out mice on 129S2/SvPas (129) (55, 56) or C57BL/6J (B6) (23, 29, 34) inbred backgrounds, but the differences in ZIKV susceptibility between these two strains have not been reported and remain unclear due to heterogeneous experimental conditions between studies. We compared the susceptibility of age-matched 129S2/SvPas-*Ifnar1^-/-^* (129-*Ifnar1*) and C57BL/6J-*Ifnar1^-/-^* (B6-*Ifnar1*) mice infected intraperitoneally (IP) with 10^7^ FFUs of FG15 ZIKV. B6-*Ifnar1* mice showed increasingly severe symptoms, with body weight loss, ruffled fur, ataxia and hind limb paralysis from day 4 p.i. and were all (10/10) moribund or dead by day 7 p.i.. By contrast, 129-*Ifnar1* mice developed mild symptoms (ruffled fur, hunched back) from day 6 p.i. with only one mouse dying on day 9 p.i. while the others recovered (FIG 1), demonstrating that the susceptibility to ZIKV infection conferred by *Ifnar1* genetic inactivation is critically influenced by the host genetic background.

**FIG 1.**
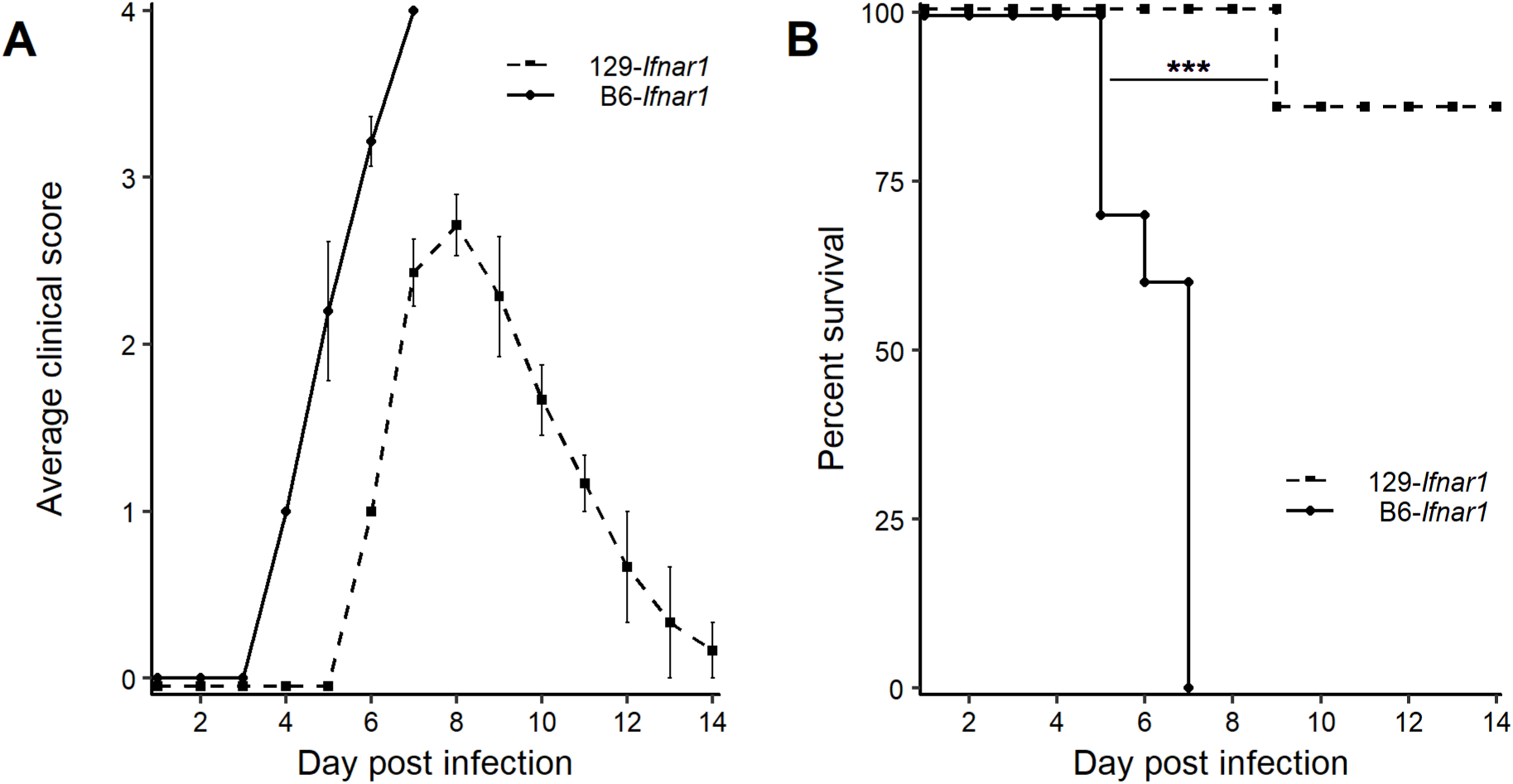
ZIKV disease severity in *Ifnar1*-deficient mice is driven by the genetic background. 6-7 week-old 129-*Ifnar1* (n = 7) and B6-*Ifnar1* (n = 10) mice were infected IP with 10^7^ FFUs of ZIKV FG15 and monitored for 14 days. (A) Average clinical score, with numerical values given as follows: 0, no symptom; 1, ruffled fur; 2, emaciation, hunched posture and/or hypo activity; 3, hind limb weakness, prostration and/or closed eyes; and 4, moribund or dead. (B) Kaplan-Meier survival curves showing 100% lethality in B6-*Ifnar1* mice at day 7 p.i. and survival of 6/7 129-*Ifnar1* mice (logrank test, ***: p = 0.0002). B6-*Ifnar1* mice developed early symptoms which rapidly evolved to death, while 129-*Ifnar1* mice developed symptoms two days later which eventually resolved in most mice.

### mAb blockade of IFNAR is a robust model to study ZIKV infection in CC mice

*Ifnar1* genetic deficiency abrogates permanently IFN-α/β-mediated immune responses but is not currently available on diverse genetic backgrounds. We therefore tested the suitability of transient IFNAR blockade mediated by mAb treatment as a model to study ZIKV infection in genetically diverse mice like the CC. Since MAR1-5A3 mAb was generated in a laboratory strain (129-*Ifnar1* mice) (30), we first assessed its efficacy by Western Blot analysis on mouse embryonic fibroblasts (MEFs) isolated from two CC strains (CC001 and CC071) both of which inherited the *Ifnar1* allele from the CAST/Ei wild-derived strain (57), by comparison with B6 MEFs. IFNAR stimulation by IFN-α/β activates the JAK1/TYK2 pathway and results in the phosphorylation of STAT1. We found that, in B6, CC001 and CC071 MEFs, STAT1 phosphorylation was equally induced by murine IFN-α and fully inhibited by the MAR1-5A3 mAb (FIG 2A).

**FIG 2.**
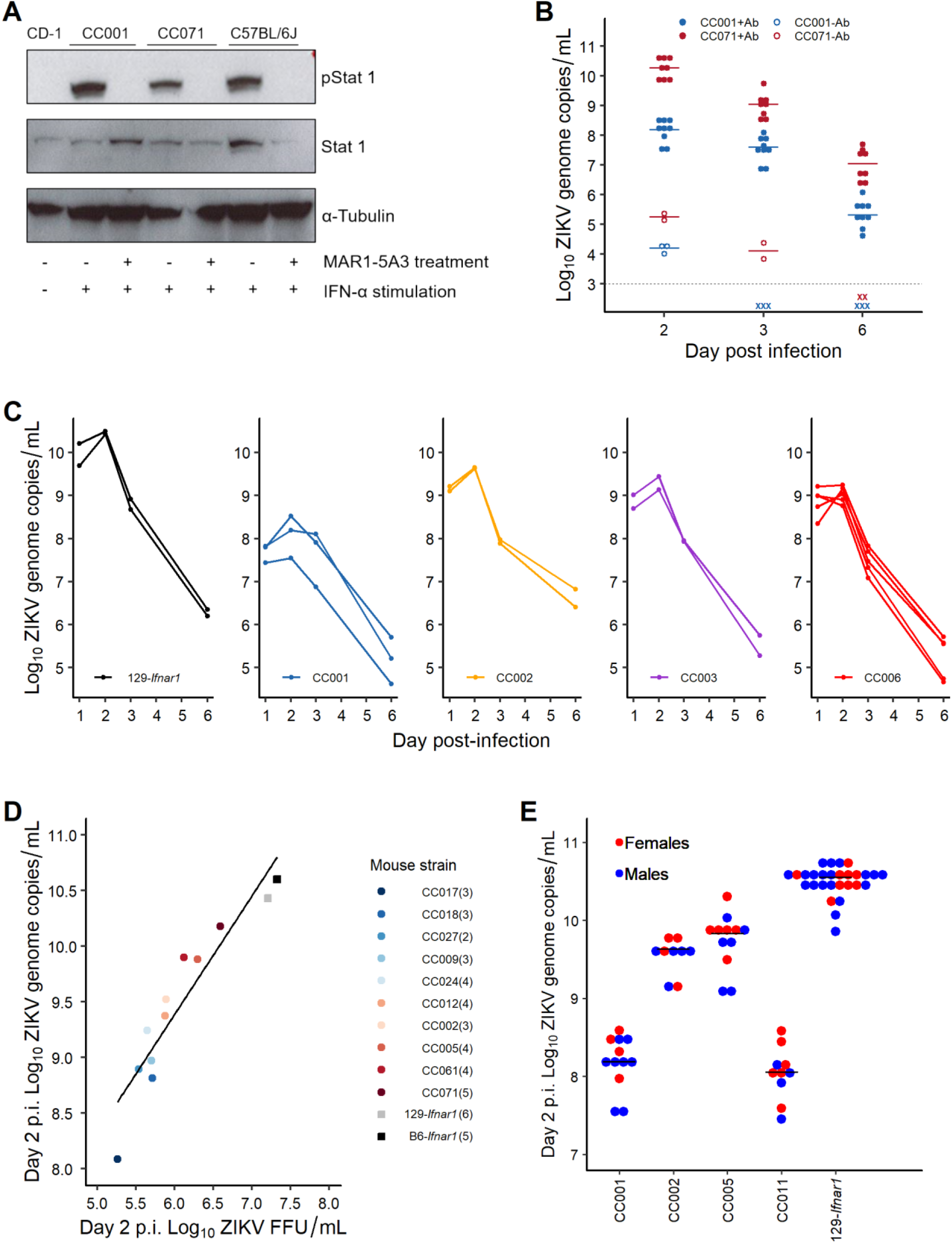
Establishment and validation of the experimental conditions for assessing susceptibility to ZIKV in CC strains. (A) The efficacy of the MAR1-5A3 mAb to block the IFNAR receptor in diverse mouse genetic backgrounds was determined by Western blotting on mouse embryonic fibroblasts (MEFs) derived from C57BL/6J, CC001 and CC071 strains. The phosphorylation of STAT1 was equally induced by IFN-α stimulation and fully inhibited by the MAR1-5A3 mAb in the three strains, as in the untreated and unstimulated CD-1 MEFs. (B) When treated with MAR1-5A3 24h prior to ZIKV infection (filled circles), mice of both CC001 and CC071 strains (n = 9 and 8, respectively) were much more permissive to viral replication with 4 to 5 log_10_ times more copies of viral genome than mice without treatment (open circles; n = 3 and 2, respectively) throughout the first three days p.i. (“x” denotes a sample below the detection level). (C) The kinetics of plasma viral load in 129-*Ifnar1* and 4 CC strains showed a maximum in most individuals at day 2 p.i. which was subsequently selected to measure peak viral load in all CC strains. Each circle represents a mouse analyzed on days 1, 2, 3 and 6. (D) Correlation between plasma viral load determined by FFA (x-axis) and RT-qPCR (y-axis) was established on 46 blood samples from 129-*Ifnar1*, B6-*Ifnar1* and 10 CC strains (circles show the mean of each strain; the number of mice per strain is shown in parentheses). The two variables were strongly correlated over a 3 log_10_ range of viral genome copies (r² = 0.89, p = 9.9x10^-17^), the number of genome copies by RT-qPCR being on average 3 log_10_ units higher than the viral titer by FFA. (E) Plasma viral load at day 2 p.i. was not significantly different (p = 0.24) between males and females of 129-*Ifnar1* and 4 CC strains for which age-matched mice of both sexes had been tested (with n ≥ 4 mice per group).

To assess MAR1-5A3 mAb efficacy *in vivo*, we infected CC001 and CC071 strains with 10^7^ FFUs of FG15 ZIKV IP and we measured the kinetics of plasma viral load in mice with and without 2 mg mAb treatment 24 hours prior to infection. Consistent with previous studies in B6 (29, Smith, 2017 #31, Scott, 2018 #25) and BALB/c (58) mice, viral load was consistently 4 to 5 log_10_ units higher in mAb-treated mice of both CC001 and CC071 strains compared to untreated mice, demonstrating that MAR1-5A3 mAb treatment successfully increases CC mice permissiveness to ZIKV replication (FIG 2B).

We then measured the kinetics of plasma viral load in 129-*Ifnar1* strain as well as in four mAb-treated CC strains infected with 10^7^ FFUs of FG15 ZIKV IP. We established that the peak plasma viral load occurred in most individuals at day 2 p.i., independently of mouse genetic background (FIG 2C).

In previous studies, viral loads have been measured either by FFU titration or by RT-qPCR quantification of viral genome copies. We compared these two methods in B6-*Ifnar1*, 129-*Ifnar1*, and in ten mAb-treated CC strains. We performed Focus Forming Assays (FFA) to measure viral particles in the plasma at day 2 p.i. and confirmed the production of infectious ZIKV in the blood of all strains (FIG 2D). Next, we compared the plasma viral load measured by RT-qPCR and by FFA. We found that these two parameters were strongly correlated over a 2 log_10_ range (Pearson coefficient, r^2^=0.89, p=9.9x10^-17^), with the number of genome copies being on average 3 log_10_ units higher than the number of FFUs (FIG 2D). We therefore validated that RT-qPCR measurement of plasma viral load could be used as a labor-efficient proxy for viremia throughout the study.

Finally, we compared plasma viral load at day 2 p.i. between males and females in 129-*Ifnar1* and in four mAb-treated CC strains in which both sexes had been tested. We found no significant difference between sexes across diverse genetic backgrounds (two-way ANOVA, p=0.24; FIG 2E), validating the use of merged data from males and females in mouse ZIKV infection experiments.

### CC genetic diversity drives ZIKV disease severity and plasma viral load

To explore broad genetic variation, we assessed the susceptibility of mAb-treated mice of 35 CC strains. B6-*Ifnar1*, 129-*Ifnar1* and mAb-treated B6 mice were included as reference strains. Only mice from three CC strains developed symptoms as shown on FIG 3A which summarizes clinical observations at day 7 p.i. CC021 and CC026 mice recovered and survived, while symptoms worsened in 7/9 (78%) CC071 mice which were moribund or died between days 7 and 9 p.i.

**FIG 3.**
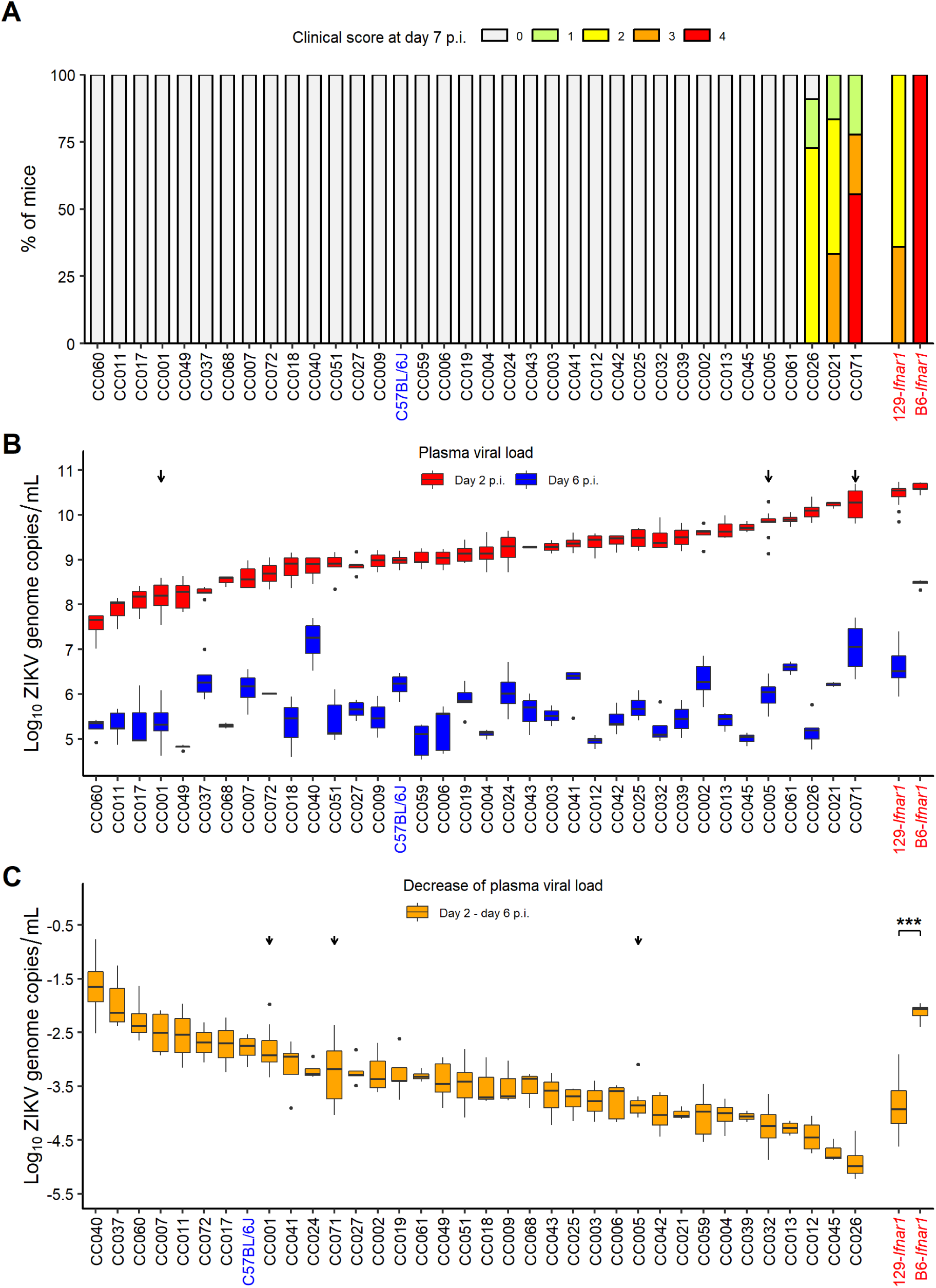
CC genetic diversity strongly impacts clinical severity and plasma viral load. Thirty-five CC strains (n = 2 to 9 per strain) were infected IP with 10^7^ FFUs of ZIKV FG15, 24hr after IP injection of 2 mg of MAR1-5A3 mAb. 129-*Ifnar1* (n = 24) and B6-*Ifnar1* (n = 5) mice were similarly infected without mAb treatment. (A) Clinical scores at day 7 p.i. as the percentage of mice in the five levels of severity (same as in FIG 1). Most CC strains did not show any symptoms, while 78% (7/9) of CC071 died before day 8 p.i. (B) Plasma viral load at days 2 (upper values) and 6 p.i. (lower values) quantified by RT-qPCR, shown as box-whisker plot with outliers as dots (strains are shown in the same order as in A). CC genetic background had a highly significant effect on viral load at day 2 p.i. (Kruskal-Wallis, p = 4.8x10^-15^) and day 6 p.i. (Kruskal-Wallis, p = 1.1x10^-10^). (C) Difference between plasma viral loads at days 2 and 6 p.i. Strains are sorted by increasing absolute difference, therefore in a different order from A and B. CC genetic background had a highly significant effect on viral load decrease (Kruskal-Wallis, p = 2.2x10^-12^). Likewise, viral load decreased much faster in 129-*Ifnar1* than in B6-*Ifnar1* mice (Wilcoxon, *** : p = 1.7x10^-5^). (B and C) Arrows indicate the subset of CC mouse strains selected for detailed study.

Plasma viral load was measured on days 2 and 6 p.i. At day 2 p.i., which corresponds to its peak, viral load was generally characterized by small within-strain heterogeneity and large inter-strain variations spread over a 2.8 log_10_ range (FIG 3B), demonstrating a strong effect of host genes (Kruskal-Wallis, p=4.8x10^-15^) with a broad sense heritability of 86% (59). The three symptomatic CC strains showed the highest peak viral load, close to that of B6-*Ifnar1* and 129-*Ifnar1* mice. However, other strains (such as CC005 and CC061) had similarly high viral loads but never showed any clinical signs of disease, indicating that peak viral load is unlikely the sole factor controlling clinical severity. At day 6 p.i., within-strain variations were larger and more heterogeneous, but we still observed highly significant inter-strain differences (Kruskal-Wallis, p=1.1x10^-10^). Interestingly, viral load on days 2 and 6 p.i. were only moderately correlated (Pearson coefficient, r²=0.46; p=0.004), indicating that viral load at day 2 p.i. was not predictive of viral load at day 6 p.i. (see for example CC018 and CC040, or CC026 and CC071).

We used the difference of the log_10_ plasma viral loads between days 2 and 6 p.i. to estimate the clearance rate of the virus from the blood stream (FIG 3C; strains sorted by increasing clearance rate, therefore differently from FIG 3A and B). This rate varied over a 3.3 log_10_ range between strains, demonstrating a strong effect of host genes (Kruskal-Wallis, p=2.2x10^-12^) with a broad sense heritability of 76%. Likewise, B6-*Ifnar1* mice showed a slower decrease in viral load than 129-*Ifnar1* mice (Wilcoxon, p=1.7x10^-5^), despite similar peak viral load at day 2 p.i.

Overall, genetic diversity in the CC panel controlled clinical severity of ZIKV infection, mouse survival, and the peak and clearance rate of plasma viral load. Of note, there was no association, across the 35 CC strains tested, between the peak plasma viral load and the *Ifnar1* allele inherited from the founder strain (ANOVA, p>0.09). This analysis confirms our *in vitro* data (FIG 2A) and indicates that the variations in peak plasma viral load do not result from differences in mAb treatment efficacy due to the *Ifnar1* alleles.

From this screening, we identified several strains with extreme phenotypes, in particular CC071 which was the most susceptible to ZIKV infection, CC001, CC011, CC017 or CC060 with low peak plasma viral load, CC040 with slowly decreasing plasma viral load and CC045 or CC026 with high peak but fast-decreasing plasma viral loads.

### CC mice show correlated susceptibility to ZIKV, DENV and WNV

We further characterized three CC strains (indicated by arrows on FIG 3B and FIG 3C) among those showing lowest (CC001) and highest peak viral loads with (CC071) or without (CC005) clinical symptoms. To establish whether the above differences were specific of the FG15 ZIKV strain of the Asian lineage, we first assessed the susceptibility of the three selected strains to the HD78788 ZIKV strain of the African lineage. 129-*Ifnar1* mice and mAb-treated CC mice were infected with 10^3^ FFUs of HD78788 ZIKV IP which proved to be highly pathogenic in *Ifnar1*-deficient mice with rapid and severe symptoms and 100% mortality (FIG 4A). CC001 was fully resistant with no or mild clinical signs (FIG 4A, left and center panels). By contrast, all CC071 mice were moribund or dead by day 10 p.i., with early and quickly aggravating symptoms, almost like 129-*Ifnar1* mice. Only 1/5 CC005 mouse died with late symptoms. Peak viral load (day 2 p.i.) varied over a 2.4 log_10_ range and the differences between strains were similar to those observed with the FG15 ZIKV strain (FIG 3B). Here again, plasma viral load at day 2 p.i. was the highest in very susceptible CC071 and 129-*Ifnar1* strains and low in resistant CC001, but it was also very high in CC005 which was moderately susceptible, confirming that clinical severity does not depend solely on peak plasma viral load.

**FIG 4.**
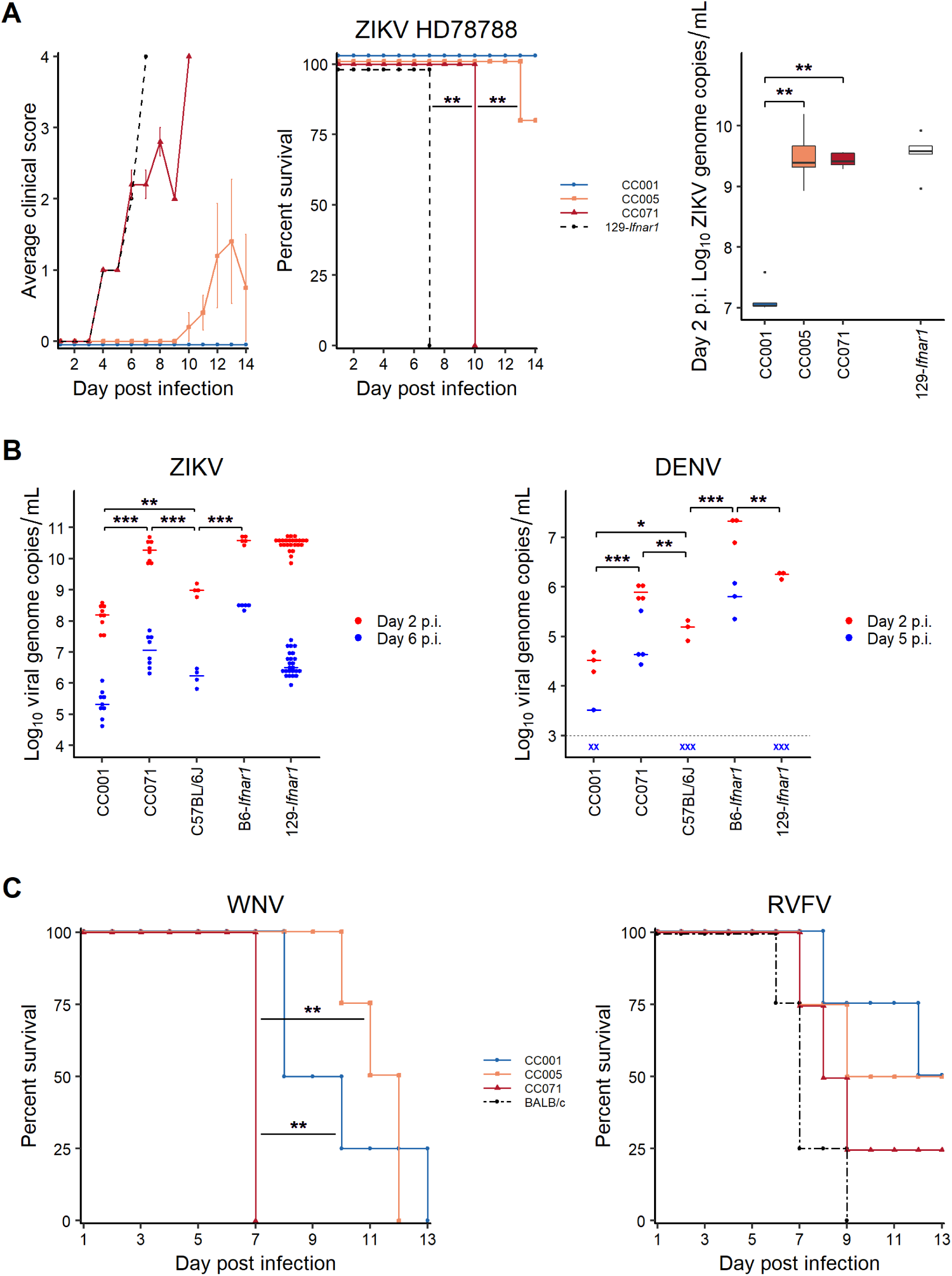
The differences in susceptibility to ZIKV between CC strains are conserved with other flaviviruses. (A) Mice from three selected CC strains treated with MAR1-5A3 mAb and 129-*Ifnar1* mice were infected intraperitoneally with 10^3^ FFUs of ZIKV HD78788. Left : average clinical score, with numerical values given as in FIG 1. CC071 and 129-*Ifnar1* mice rapidly developed severe symptoms and died while CC001 and CC005 mice were mostly resistant. Center : Kaplan-Meier survival curves (logrank test). Right : plasma viral load at day 2 p.i., measured by RT-qPCR and expressed as ZIKV genome copies per milliliter, was much lower in CC001 than in the two other CC strains (Wilcoxon). (B) Viral load after ZIKV infection (left, data extracted from FIG 3) and DENV infection (right, IV infection with 2.10^6^ FFUs of DENV KDH0026A) was compared in mAb-treated CC001, CC071 and B6 mice and in 129-*Ifnar1* and B6-*Ifnar1*. Most between-strain differences were conserved between the two viruses (*t* test). (C) Left : Kaplan-Meier survival curves of four male mice of each of the three selected CC strains infected IP with 1000 FFU of WNV strain IS-98-ST1 and monitored for 14 days. CC071 died earlier than CC005 and CC001 mice (logrank test). Right : Kaplan-Meier survival curves of four to five male mice of BALB/cByJ and each of the three selected CC strains infected IP with100 PFUs of RVFV strain ZH548 and monitored for 14 days. No significant difference was found among the three CC strains and only CC001 mice survived longer than BALB/c mice (logrank test, p > 0.05). * p < 0.05; ** p < 0.01; *** p < 0.001.

To evaluate whether these differences in susceptibility were specific to ZIKV or extended to other flaviviruses, we assessed the phenotype of a few strains after infection with DENV and WNV, two other members of the *Flaviridae* family.

We measured plasma viral load after IV infection with 2x10^6^ FFUs of KDH0026A DENV in mAb-treated CC001, CC071 and B6 mice and in 129-*Ifnar1* and B6-*Ifnar1* mice (FIG 4B right). Most inter-strain differences observed with ZIKV FG15 strain (FIG 4B left, data from FIG 3B) were conserved with DENV, CC071 displaying the highest plasma viral load in Ab-treated mice. DENV infection was overall much less clinically severe since only B6-*Ifnar1* mice developed non-lethal symptoms including ruffled fur, hunched back and ataxia.

We also investigated the susceptibility of the selected CC strains to WNV. *Oas1b* was previously shown to be a major host genetic determinant of susceptibility to WNV in mice (60). Of note, the three selected CC strains carry the same non-functional allele of *Oas1b* inherited from the laboratory strain founders, conferring them susceptibility to WNV infection. CC mice were infected IP with 10^4^ FFUs of WNV IS-98-ST1 and monitored for 14 days p.i. (WNV infection does not require anti-IFNAR mAb treatment in *Oas1b*-deficient mice). All CC071 mice died 7 days p.i., significantly faster than CC001 and CC005 mice (logrank, p<0.01; FIG 4C, left panel), indicating that genetic diversity between CC strains also influences their susceptibility to WNV even in the context of *Oas1b* deficiency.

To assess whether the differences of susceptibility between these CC strains also applied to other viruses, we infected them with 10^2^ PFUs of RVFV ZH548 IP. No significant difference was found between CC strains (logrank, p>0.05 for all pair comparisons; FIG 4C, right panel) which succumbed late from the infection, like the commonly used BALB/cByJ mice.

### Genetic analysis suggests a polygenic control of susceptibility to ZIKV in CC mice

To identify host genetic factors controlling the susceptibility to ZIKV in CC strains, we performed a genome-wide association study between the plasma viral loads at days 2 and 6 p.i. or the decrease rate of plasma viral load, and the genotypes of the 35 CC strains. Genetic associations were plotted as LOD scores (FIG 5). We did no find genome locations at which LOD scores reached the minimum 0.1 significance threshold for any of the three traits, while it would be expected if phenotypic variations were controlled by one or two loci with strong effects. Therefore, these results suggest that plasma viral load is controlled by multiple small-effect genetic variants.

**FIG 5.**
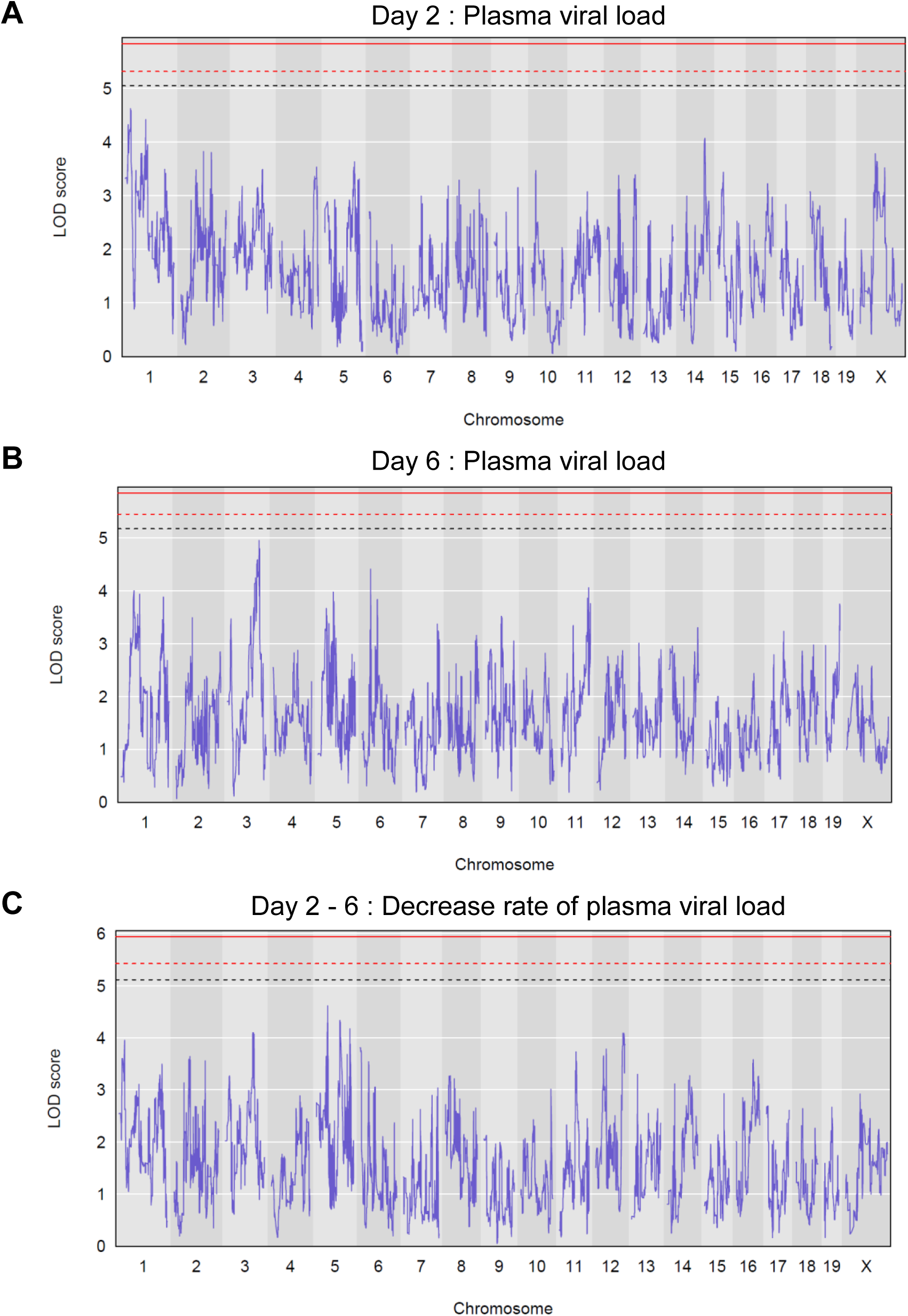
Genetic analysis of susceptibility to ZIKV fails to identify simple genetic control. Genome-wide linkage analysis for the plasma viral load at day 2 p.i.(A), the plasma viral load at day 6 p.i. (B) and the decrease rate of plasma viral load (C) on the 35 CC strains shown on FIG 3. The x-axis represents genomic location; the y-axis is the LOD score, representing the statistical association between the phenotype and the genomic location. Genome-wide thresholds p = 0.1, p=0.05 and p = 0.01, computed from 1000 permutations, are represented by dashed black, dashed red and plain red lines, respectively. No genome location reached the p = 0.05 threshold.

### Genetic diversity of CC strains controls brain viral load and pathology

To assess the influence of host genetics on the brain pathology caused by ZIKV infection, we further characterized the three previously selected CC strains. We measured the viral load in the brain 6 days after IP infection with FG15 ZIKV in mAb-treated CC mice and 129-*Ifnar1* mice (FIG 6, top). CC005 and CC071 which had higher peak plasma viral load also had higher brain viral load (FIG 6, mean=6.5 log_10_ copies/µg RNA for CC005 and CC071, compared with 5 log_10_ copies/µg RNA for CC001). As expected, 129-*Ifnar1* mice showed the highest viral load in the brain. These results indicate overall correlation between plasma and brain viral loads.

**FIG 6.**
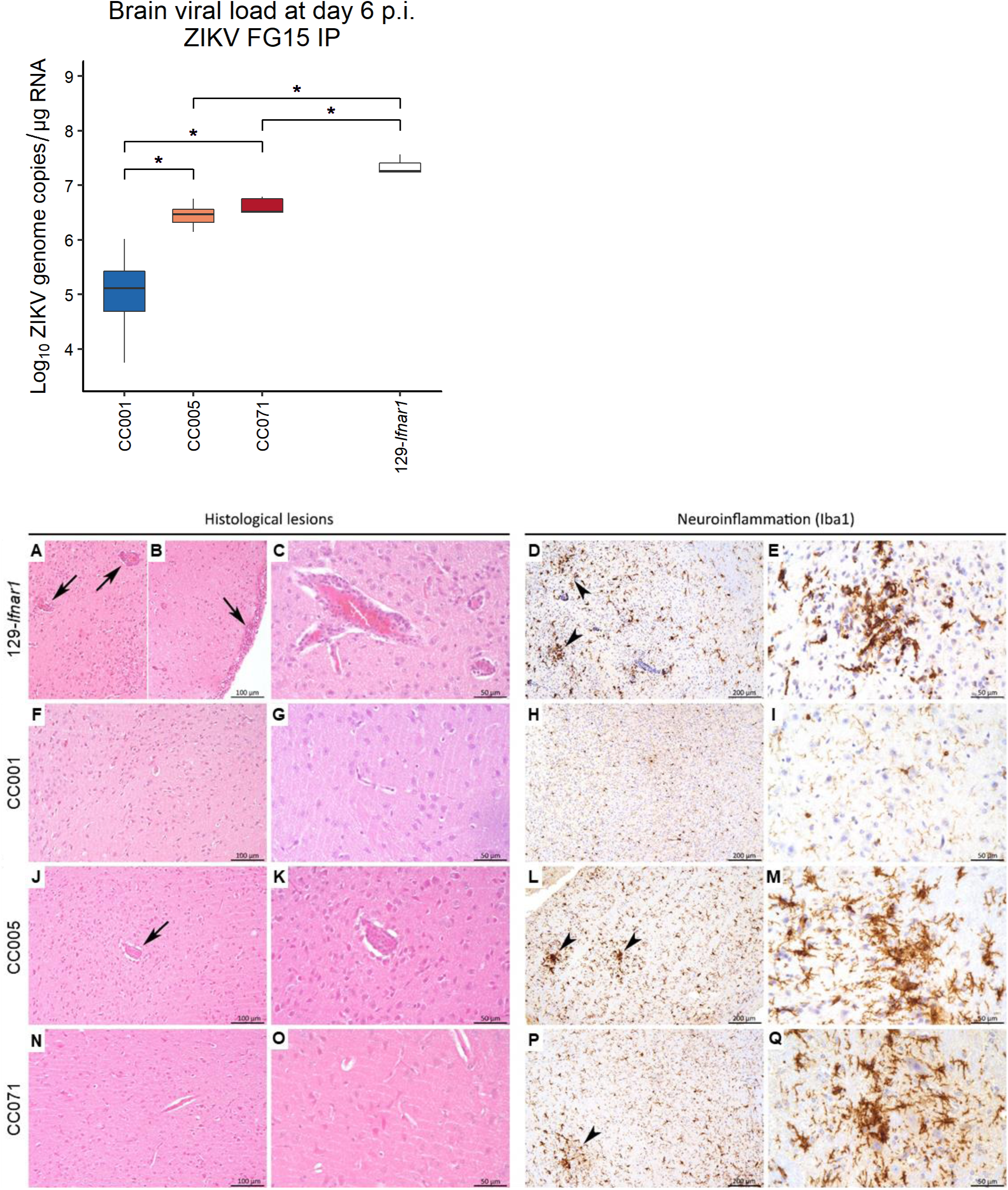
Genetic variations between CC strains control brain viral load and histological profile of infected mice. Four to five mice of 129-*Ifnar1* and three selected CC strains were infected IP with 10^7^ FFUs of ZIKV FG15 24hr after IP injection of 2 mg of MAR1-5A3 mAb. Top : Brain viral load was measured by RT-qPCR at day 6 p.i. (Wilcoxon, * : p < 0.05). Bottom : Representative brain sections of the same ZIKV FG15-infected mice, at day 6 p.i. (A-C) Lesions were clearly more severe for 129-*Ifnar1* mice (n=3), with subacute leptomeningo-encephalitis (*i.e.* infiltration of perivascular spaces and leptomeninges by lymphocytes, plasma cells and macrophages; arrows) and (D-E) activation of microglial cells with microglial nodules (arrowheads). (F-G) CC001 mice (n=5) displayed no significant histological lesions in the brain with normal resting (non-activated) microglial cells (H-I). Only very rare small clusters of activated microglial cells were detected (data not shown). By contrast, CC005 mice (n=5) displayed moderate inflammatory lesions characterized by (J-K) perivascular cuffing (arrow; n=2), (L-M) activation of microglial cells (hyperplasia and thickening of cell processes) and microglial nodules (arrowheads; n=5). CC071 mice (n=4) also displayed inflammatory lesions but with intermediate severity: (N-O) almost no lesions in HE but (P-Q) activation of microglial cells and microglial nodules (arrowhead). A, B, C, F, G, J, K, N, O : HE staining; D, E, H, I, L, M, P, Q : anti-Iba1 immunohistochemistry.

Histopathological analysis, carried out in the brain of the same mice (FIG 6 bottom), revealed different lesion profiles between the four mouse strains. 129-*Ifnar1* mice indeed displayed the most severe inflammatory lesions (subacute leptomeningo-encephalitis). By contrast, almost no lesions were detected in the brain of CC001 mice. CC005, and CC071 displayed only minimal to mild encephalitis (more severe for CC005 mice), but with activation of microglial cells and microglial nodules similar to that of 129-*Ifnar1* mice, as revealed by Iba1 immunolabeling.

The nature and intensity of brain histological lesions may depend on the circulating viral load, on the capacity of the virus and of the mAb to cross the blood-brain barrier and on the permissiveness of brain cells (in particular neurons and microglia). To assess the differences in susceptibility of brain cells between CC strains, we performed intra-cerebral infections to deliver the virus directly into the brain tissue. 129-*Ifnar1* and mAb-untreated CC mice received 10^5^ FFUs of FG15 ZIKV in the left ventricular region of the brain and were followed for 3 weeks. Mild and transient symptoms (ruffled fur, hunched back) were observed in a few mice of the three strains and one CC005 mouse died on day 19 p.i. A second group of CC mice were infected similarly and euthanized at day 6 p.i. for histological analysis. Differences in brain viral load between CC strains were similar to those observed after IP infection, with CC005 and CC071 mice showing significantly higher brain viral load than CC001 mice (FIG 7, top). Compared with IP route of infection, lesions in 129-*Ifnar1* mice were mostly similar, while lesion profiles were clearly different in the three CC strains (FIG 7, bottom). Strikingly, the most severe lesions were detected for CC071, with marked subacute leptomeningo-encephalitis and strong activation of microglial cells (Iba1 staining). CC001 and CC005 mice also displayed inflammatory lesions, clearly less severe than CC071, with gliosis, microglia activation and microglial nodules.

**FIG 7.**
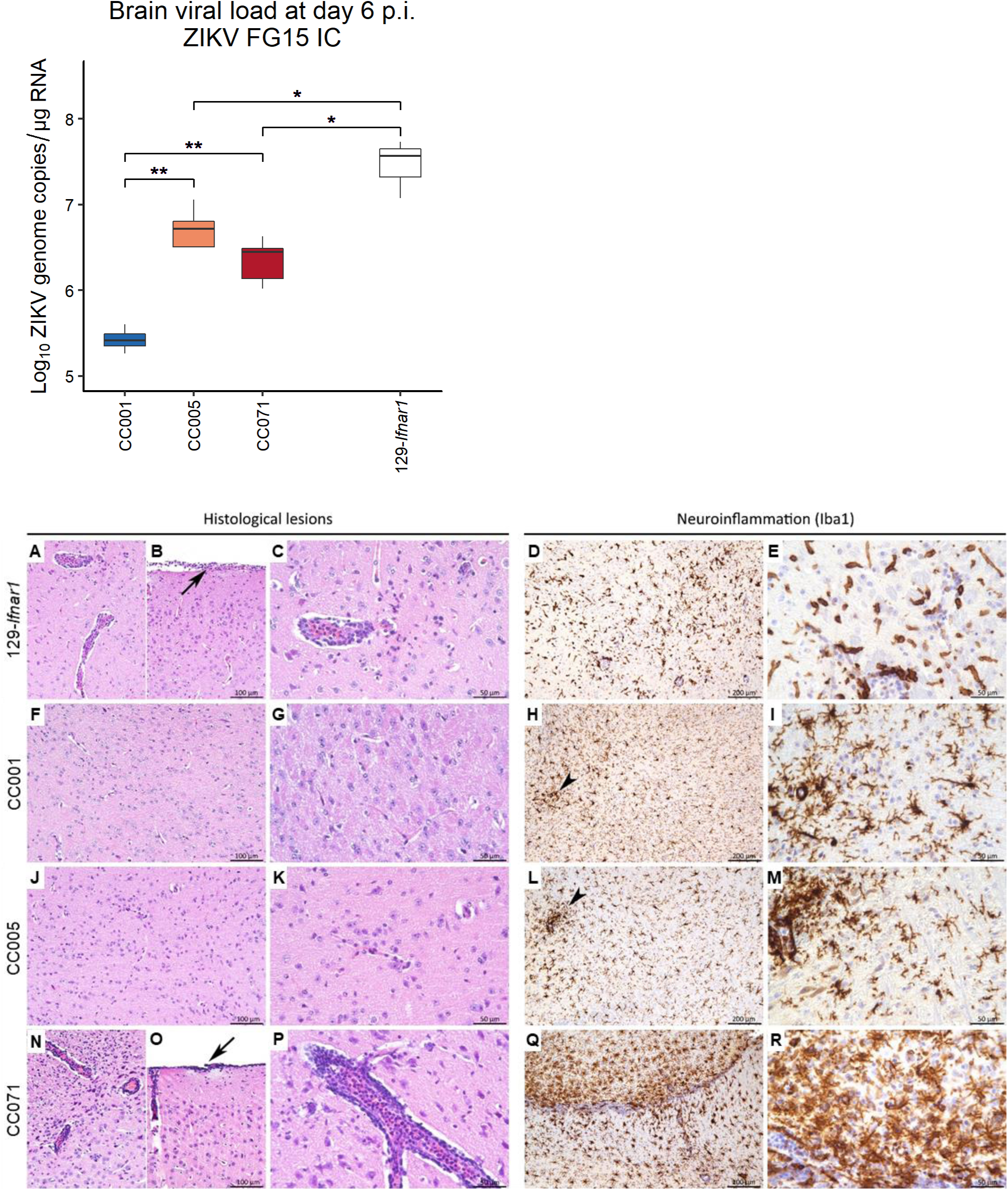
Intracranial ZIKV FG15 infection results in strain-dependent viral load and brain histological lesions. Mice of 129-*Ifnar1* and three selected CC strains (3-5 mice per strain) were infected IC with 10^5^ FFUs of ZIKV FG15 in the absence of prior anti-IFNAR treatment. Top: CC005 and CC071 mice show significantly higher brain viral load at day 6 p.i. than CC001 mice (Wilcoxon, ** : p < 0.01). Bottom : After intracranial inoculation, lesion profiles were clearly different from those observed after IP inoculation. (A-C) 129-*Ifnar1* mice (n=4), still displayed marked subacute leptomeningo-encephalitis (arrow: leptomeningitis) and (D-E) activation of microglial cells. Two of the 5 CC001 mice displayed no significant histological lesions with normal resting microglial cells, while the other three displayed (J-K) minimal lesions with gliosis, and (H-I) rare small clusters of activated microglial cells. In this experimental model, CC005 mice displayed heterogeneous lesion profiles with either (i) suspected meningitis and (J-K) gliosis (n=4/5), or (ii) moderate leptomeningo-encephalitis (n=1/5). (L-M) Activation of microglial cells (arrowhead), with variable severity, was detected in all animals (n=5/5). Strikingly, (N-P) all CC071 mice (n=5/5) displayed marked leptomeningo-encephalitis (arrow: leptomeningitis) with (Q-R) strong activation of microglial cells. A, B, C, F, G, J, K, N, O, P : HE staining; D, E, H, I, L, M, Q, R : anti-Iba1 immunohistochemistry.

These results indicate that CC strains differ in their permissiveness to viral replication in the brain and in their susceptibility to ZIKV-induced histological brain damage, independently from potential differences in the capacity of ZIKV to disseminate to the brain from the circulation.

### Viral replication in CC071 cells is increased *in vitro*

Differences in peak plasma viral load and results from IC infections suggested that different rates of viral replication could contribute to the variations in susceptibility between CC strains. To address this point, we measured the production of viral particles in three cell types infected with ZIKV FG15. We derived primary MEFs, peritoneal macrophages (PMs) and primary cultured neurons (PCNs) from CC001 and CC071 strains. Cells were infected with ZIKV FG15 at a MOI of 5. In all three cell types, CC071 cells produced increasingly higher amounts of viral infectious particles than CC001 cells between 24 and 72 hours (FIG 8). These results suggest that increased replication rate in CC071 could contribute to its susceptible phenotype.

**FIG 8.**
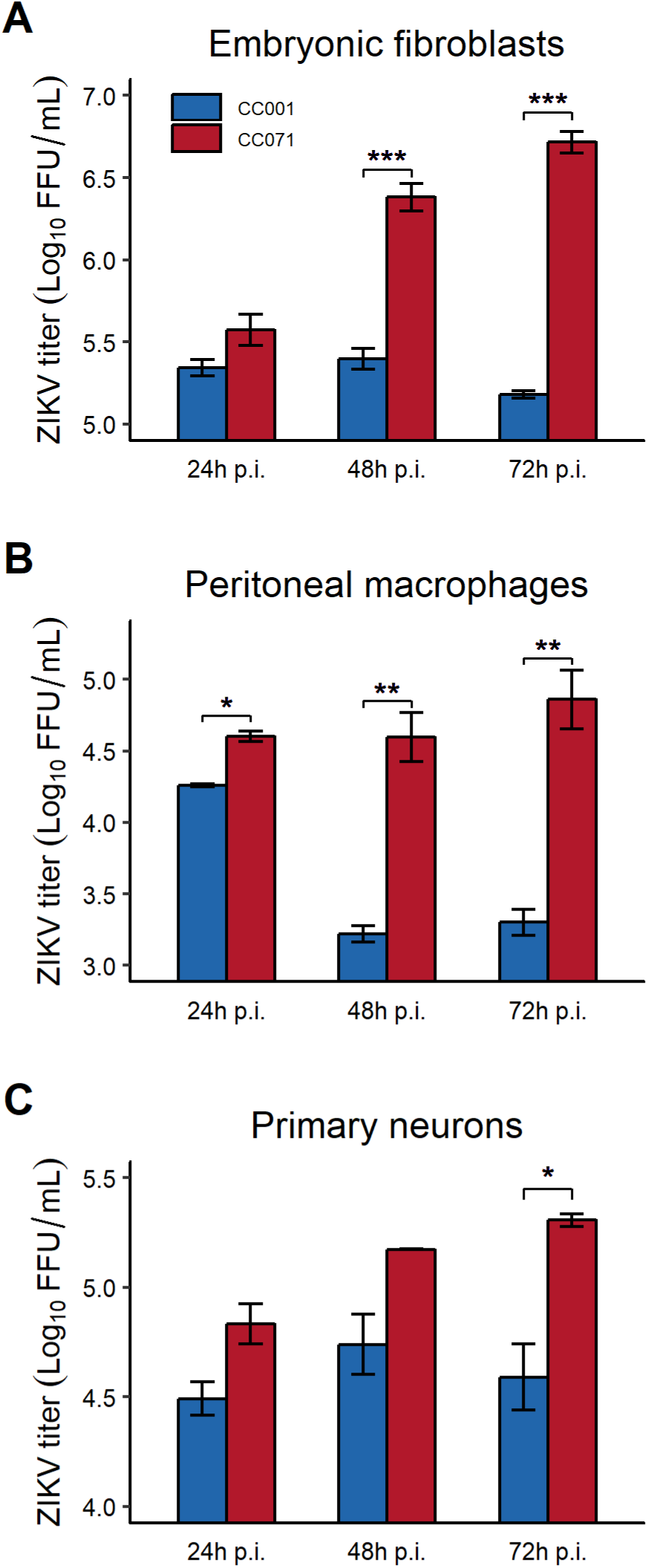
*In vivo* susceptibility to ZIKV FG15 in CC071 correlates with increased viral replication *in vitro* compared with CC001. Mouse cells were infected with ZIKV FG15 at MOI 5. ZIKV titer in the supernatant was quantified by focus-forming assay at 24, 48 and 72 hours p.i. (A) MEFs derived from CC001 (blue) and CC071 (red) embryos. Mean +/- SEM from 3 biological replicates. At 48h and 72 h p.i., CC071 MEFs produced significantly higher virus titers. (B) Macrophages isolated from peritoneal lavage. Mean +/- SEM from 2 replicates. CC071 macrophages produced significantly higher virus titers at the three time points. (C) Neurons from fetal brain dissected at day 16.5 of gestation and cultured for 12 days before infection. Mean +/- SEM from 2 replicates. CC071 neurons produced significantly higher virus titers at 72 h p.i. (t tests; * p < 0.05, ** p < 0.01, *** p < 0.001).

## DISCUSSION

ZIKV is a serious public health concern considering the occurrence of severe neurological complications in adults and congenital malformations that can result from the infection of pregnant women. The variable outcomes of ZIKV infection in humans has led to hypothesize a role for host genetic factors (9, 13) although this has never been demonstrated thus far. As for other infectious diseases, human genetic studies on susceptibility to ZIKV would require large cohorts of patients and would be confounded by pathogen genetics, pathogen dose, mosquito-dependent factors and multiple environmental parameters.

Several mouse models of human ZIKV infection have already been described and have substantially improved our understanding of viral tropism, dissemination, pathogenesis, persistence, transmission and vaccine protection. To overcome the inability of ZIKV to inhibit in mice IFN induction and signaling pathways as observed in humans (27), most studies have been performed using *Ifnar1*-deficient mice which have become a reference model. However, high levels of viral replication can also be achieved by temporary inhibition of IFN signaling by anti-IFNAR mAb treatment (30, 31, 61) or even in immunocompetent mice by infecting neonates (36, 62–64) or using a combination of mouse-adapted ZIKV strains and human STAT2 knock-in mice (65).

The choice of the ZIKV strain used in an animal model is important to maximize the relevance of mouse studies to human infection. Mouse studies have used different ZIKV strains from the African or Asian lineages. Mouse-adapted strains of the African lineage derived from a large number of serial passages are more pathogenic in mice at lower doses (34) but carry mutations that may bias the translatability of results to humans. To avoid this limitation, mouse studies have often used different ZIKV strains from the Asian lineage derived from clinical isolates. Genetic differences between these two lineages are suspected to be responsible for the emergence of symptomatic cases in human starting with the Yap Island epidemics in 2007 (7, 8). Therefore, while the ZIKV strain needs to be standardized in experimental studies, generalization of the results obtained with one viral strain require confirmation using another strain. Because of the incidence of neurological complications associated with infections by Asian lineage ZIKV, we used for our genetic screening a low-passage strain derived from a 2015 case of French Guyana, at an early stage of the South-American epidemics. Since this strain had not been adapted to the mouse, high doses were required to achieve high circulating viral loads.

Most mouse studies have used either B6-*Ifnar1* or 129-*Ifnar1* strains without specific rationale and their results cannot be directly compared due to many experimental differences such as ZIKV strain, dose and route of inoculation (66). Under strictly identical conditions, we found that B6-*Ifnar1* mice developed more rapid and severe clinical symptoms and higher mortality than 129-*Ifnar1* mice, despite similar levels of plasma viral RNA at day 2 p.i. We also found that viral load persisted longer in B6-*Ifnar1* mice. These results show that, under our experimental conditions, these two *Ifnar1*-deficient strains have clearly distinct susceptibility to ZIKV. To our knowledge, these two strains have been compared in only one study which found no difference in survival after WNV infection (67). However, their extreme susceptibility might have prevented the identification of any difference. Our results have practical implications for many studies based on *Ifnar1*-deficient mice and motivate further genetic studies to identify the determinants and mechanisms controlling differences of susceptibility between B6 and 129 inbred backgrounds.

To further investigate the role of host natural genetic variants on ZIKV susceptibility, we leveraged the genetic diversity across CC strains. The CC has been developed as a collection of inbred strains that more accurately reproduce the genetic diversity and phenotypic range seen in human population (68). To enable systemic ZIKV replication after parenteral inoculation in diverse genetic backgrounds, we blocked type I IFN response using MAR1-5A3 mAb (23, 31). However, because CC genetic diversity includes sequence polymorphisms in the *Ifnar1* gene which could affect the efficacy of mAb-mediated IFNAR inhibition, we confirmed full abrogation of IFNα-induced STAT1 phosphorylation in MEFs from two CC strains carrying a wild-derived *Ifnar1* haplotype. Moreover, we showed that the differences in peak plasma viral load across 35 CC strains were not associated with the *Ifnar1* allele each CC has received from the founder strains. These results validate that the MAR1-5A3 mAb has similar efficacy across a broad range of mouse genetic backgrounds, which will be useful to develop new models of viral infections.

A single injection of MAR1-5A3 mAb 24 hours before ZIKV infection resulted in moderate to very high levels of viral RNA in the blood and brain. ZIKV infection was symptomatic in a minority of CC strains (3/35), as observed in infected humans (9, 69), and mortality was observed only in CC071. These results confirm that ZIKV can replicate and establish viremia without inducing symptoms (29). Moreover, while all symptomatic strains had high peak viral loads, other strains with similar viral loads (like CC005 or CC061) never developed any signs of illness, indicating that other pathogenic mechanisms are required to result in symptomatic infection and that viral load alone does not reliably predict clinical outcome of ZIKV infection in a genetically diverse mouse population.

Since all experimental parameters were carefully standardized between strains (in particular, the microbiological environment in which they were bred), which resulted in small intra-strain variations, and since the MAR1-5A3 mAb treatment was similarly effective across strains, differences in peak viral load between strains can be confidently attributed to host genetic variants. The 86% broad sense heritability further indicates that genetic background is the principal factor driving peak viral load across CC strains.

Viremia decreased between days 2 and 6 p.i., as previously reported in several studies (55, 70, 71) but not in others (29, 56) for reasons that have not been discussed and remain unclear. In our study, the rate of decrease, which was estimated as the difference in viral load between day 2 and day 6 p.i., showed remarkable homogeneity between individuals of the same CC strain and very large variations across CC strains. This data resulted again in high broad sense heritability which demonstrates a strong influence of host genes on this trait. The decrease of circulating viral load is the net result of ZIKV production in infected tissues, dissemination to the blood stream and elimination from the circulation. Therefore, host genes could control the kinetics of viral load through multiple mechanisms.

After exploring the range of susceptibility to ZIKV across broad genetic diversity, we focused our study on a few CC strains exhibiting contrasted phenotypes with the aim of characterizing new models (72). CC001 is one of the least permissive to ZIKV, with low peak viral load. At the other extreme of the distribution, CC005 and CC071 have similarly high plasma viral loads while only CC071 shows symptoms and high mortality. These differences between CC strains were strikingly conserved with the African, mouse-adapted, HD78788 strain (FIG 4A). The use of lower infectious doses with HD78788 virus was supported by its higher pathogenicity resulting from mouse adaptation. The consistency between these two experiments suggest that the large phenotypic diversity we have reported should apply to most ZIKV strains.

Overall, brain viral load and brain pathology after IP infection were consistent with peak plasma viral load. In CC mice, the most notable microscopic lesions included signs of neuroinflammation evidenced by Iba1 immunohistochemistry. Neuroinflammation was similar in 129-*Ifnar1*, CC005 and CC071 mice. These changes were less pronounced than in a previous study which reported more severe CNS lesions in MAR1-5A3-treated B6 mice (31), infected with a more virulent African lineage ZIKV strain. The variable severity of lesions observed in CC mice, ranging from very mild abnormalities in CC001, to inflammatory lesions with perivascular cuffing, activation of microglial cells and microglial nodules in CC005, indicates that the genetic background also controls ZIKV neuropathogenesis.

The less severe histological lesions observed in CC mice compared with 129-*Ifnar1* mice could be due to the limited access to the brain of the virus or of the mAb which does not appreciably cross the blood-brain barrier (29). Therefore, intracerebral infection aimed at comparing brain lesions between strains while controlling the amount of virus effectively delivered. Surprisingly, CC001 and CC005 mice showed similar types and severity of lesions (although not all CC001 mice showed lesions) while CC071 mice developed much more severe signs of leptomeningo-encephalitis with massive neuroinflammation, similar to those of 129-*Ifnar1* mice. This last result suggests that the milder lesions observed in CC071 compared with 129-*Ifnar1* mice after IP infection were likely due to reduced viral dissemination to the brain. Importantly, mice did not receive prior mAb treatment, allowing for the development of local and systemic antiviral responses. These results emphasize the complex interplay between infected cells and effectors of the immune response, which likely differs between CC strains under the control of host genes.

Viral replication rate between resistant CC001 and highly susceptible CC071 mice was investigated as a plausible mechanism for the differences in susceptibility between these two strains. MEFs are a semi-permanent source of cells which have been extensively used to assess viral replication (73, 74) including with ZIKV (75, 76). Although not the primary target of ZIKV infection, macrophages are also a relevant cell type to investigate ZIKV replication and innate responses (77). Mouse peritoneal macrophages are more easily recovered than MEFs (78) although they can only be used as a primary culture. Finally, primary cultured neurons are of particular relevance considering ZIKV tropism for neural progenitors (63). Although the kinetics of viral replication were different between cell types, with a swift drop in CC001 macrophages at 48h while it was slower in CC001 MEFs and primary neurons, replication steadily increased over time in all three cell types of CC071 origin, leading to significantly higher viral titers at 72h. Our data is consistent with the observation by Caires-Junior et al. who reported increased ZIKV replication rate in iPS-derived neuroprogenitor cells from CZS-affected babies compared with their unaffected dizygotic twin (13). Therefore, our results strongly suggest that increased replication rate in CC071 compared with CC001 likely contributes to its higher plasma and brain viral loads and to its higher overall susceptibility to ZIKV.

Investigating the genetic diversity of a large number of CC strains has significantly extended the range of phenotypes induced by ZIKV infection in mice and better model the heterogeneity of the human population. It has allowed testing important factors such as mouse gender and the method of viral load measurement across multiple host genetic background, providing robust conclusions (37). Importantly, we found no differences between male and female mice in their susceptibility to ZIKV disease, nor in the peak viral load (FIG 2E). We also found a high correlation between viral loads measured by titration and by qRT-PCR over a 2 log_10_ range (FIG 2D). This is in contrast with a study on Ebola virus which showed that, in spleen and liver, the susceptible mice produced similar amounts of viral genomes but 1 to 2 log_10_ more infectious virions than the resistant mice (43).

Genetic diversity also allowed us to assess correlations between traits, which cannot be achieved in a single strain. We showed that brain viral load was consistent with plasma viral load, but that plasma viral loads at days 2 and 6 p.i. were only moderately correlated. Likewise, we found that clinical severity did not correlate with the intensity of brain histological lesions and neuroinflammation, as summarized in Table 1. These dissociations between phenotypes provide evidence for partly distinct mechanisms and genetic control (37) and result in distinct mouse models.

**Table 1.**
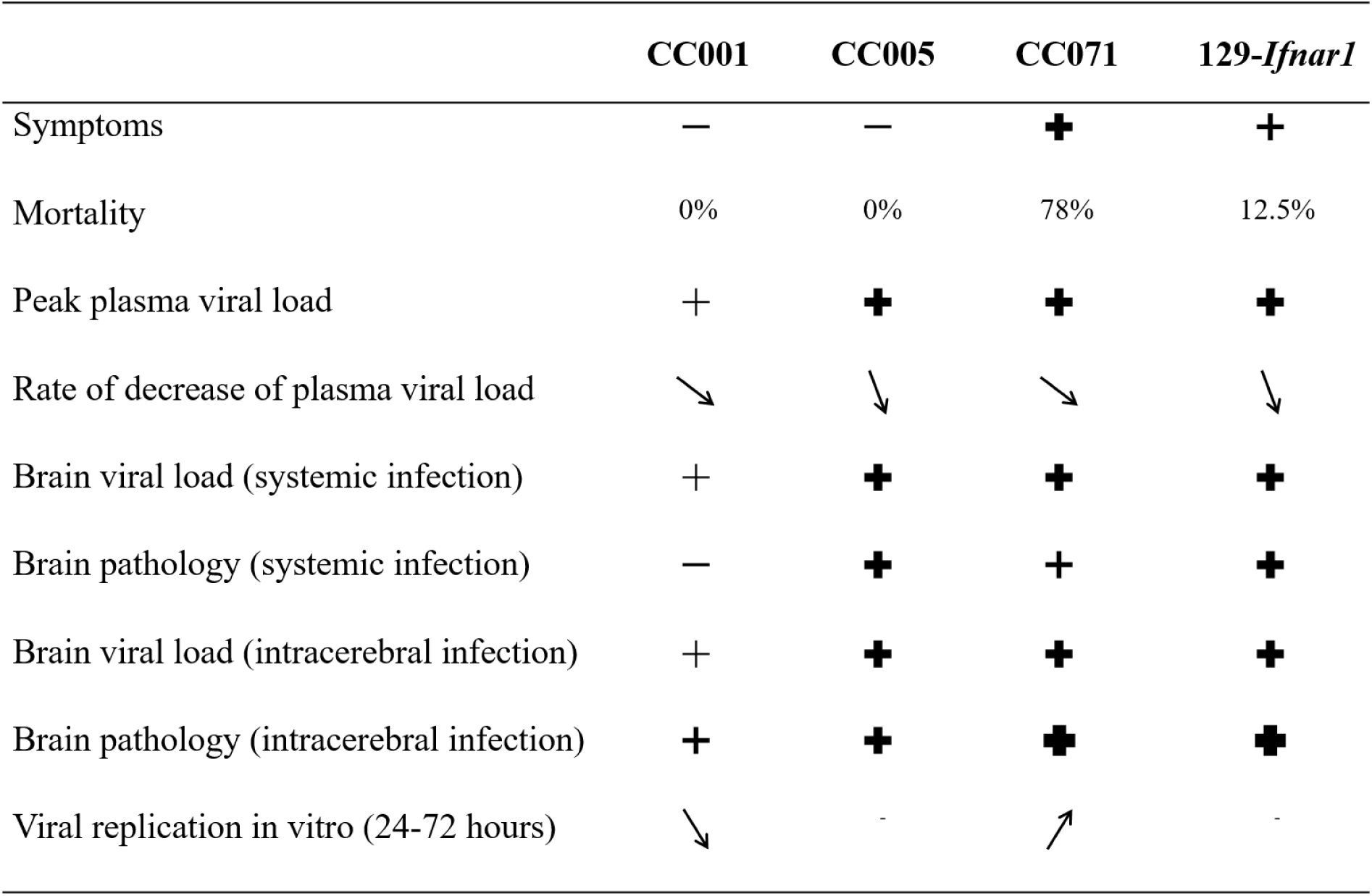
Summary of the main features of ZIKV infection in mAb-treated CC strains and 129-*Ifnar1* mice.

A recent study has reported strain-dependent variations in the long-term neuropathological and behavioral consequences of ZIKV infection after neonatal infection between four mouse inbred strains known to differ in their susceptibility to pathogens (36). Since they are all laboratory strains, they do not cover the same genetic variation as in our study and it is likely that even more diverse phenotypes would be observed in this model with the CC panel.

Genetic analysis of our results strongly suggests that, by contrast with other viruses for which major host genetic determinants have been identified (e.g. *Oas1b* for WNV (60) or *Mx1* for influenza virus (79)), susceptibility to ZIKV in CC strains is under polygenic control. This is supported by the continuous distributions of peak plasma viral load (FIG 3B) and of the rate of viral decrease (FIG 3C), and by the absence of any regions of the genome significantly associated with variations in viral load (FIG 5). Calculations based on CC genotypes show that, with 35 strains and an average of 5 mice per strain, we had 80% power of detecting a bi-allelic QTL explaining 30% or more of the phenotypic variance (80). This clearly rules out the possibility that the phenotypic variations measured across CC strains were controlled by one or few genes with major effects, as observed with *Oas1b* for WNV (47). Dissecting the genetic architecture of resistance and susceptibility to ZIKV in these strains will require dedicated intercrosses (72).

*Oas1b* is an interferon-stimulated gene and a major determinant of mouse susceptibility to WNV (45). A variant in OAS3, a member of the human homologous gene family, has been associated with increased severity of dengue (81). Most laboratory strains, including five of the eight CC founders, carry the same non-functional allele of *Oas1b* which renders them susceptible to WNV infection (60), while the three wild-derived CC founders carry polymorphic but functional alleles and are resistant (45). CC strains therefore carry either functional or non-functional *Oas1b* alleles. Our results provide multiple lines of evidence to rule out a significant role of *Oas1b* in the variations of susceptibility to ZIKV across CC strains. First, since mAb-mediated blockade of type I IFN response likely inhibits temporarily *Oas1b* induction, *Oas1b* allele is unlikely to explain differences in peak viral load at day 2 p.i. Moreover, our QTL mapping analysis showed that mouse genotype at *Oas1b* (located on distal chromosome 5) did not significantly contribute to variations in viral load at day 2 or day 6 p.i., or in the rate of viral load decrease (FIG 5). Finally, since both CC001, CC005 and CC071 strains carry the *Oas1b* deficient allele (http://csbio.unc.edu/CCstatus/CCGenomes/), their differences in clinical severity, brain pathology and replication rate in infected cells must be controlled by other genetic variants. Interestingly, the large difference of survival time after WNV infection between CC071 and CC001 or CC005 provides ideal strain combinations to identify novel genes controlling susceptibility to this virus.

Out of this large series of CC strains, we identified several new mouse models of ZIKV disease. CC071 mice were the most susceptible to ZIKV infection, more than mAb-treated B6 mice (FIG 3A,B). mAb treatment was required to achieve high circulating viral load (FIG 2B), showing that CC071 has functional type I IFN response. CC071 mice were also very susceptible to DENV and WNV, two flaviviruses related to ZIKV. However, they are not uniformly susceptible to infectious agents since they showed susceptibility to RVFV similar to that of BALB/c, CC001 and CC005 mice, and intermediate susceptibility to *Salmonella* Typhimurium (52). Together with other susceptible strains like CC021 and CC026 which developed symptoms, or CC005 which developed severe brain lesions, CC071 will help identifying mechanisms of severe ZIKV infection and their genetic control. By contrast, CC001 mice were highly resistant, even to a strongly pathogenic African ZIKV strain, despite blockade of type I IFN signaling. Extensive analysis of these CC strains with extreme phenotypes may elucidate how genetic variants affect susceptibility as well as innate and adaptive immune responses to flaviviral infection (72) and provide deeper understanding of the pathophysiology of severe complications of human ZIKV disease.

## MATERIALS AND METHODS

### Mice

All Collaborative Cross (CC) mice (purchased from the Systems Genetics Core Facility, University of North Carolina and bred at the Institut Pasteur) (82), C57BL/6J mice (purchased from Charles River Laboratories France), BALB/cByJ and *Ifnar1* knock-out mice (*Ifnar1^tm1Agt^* allele on 129S2/SvPas or C57BL/6J background, designated 129-*Ifnar1* and B6-*Ifnar1*, respectively, and bred at the Institut Pasteur) were maintained under SPF conditions with 14:10 light-dark cycle and *ad libitum* food and water in the Institut Pasteur animal facility. In all experiments, mice were killed by cervical dislocation. All experimental protocols were approved by the Institut Pasteur Ethics Committee (projects #2013-0071, #2014-0070, #2016-0013, #2016-0018 and dap190107) and authorized by the French Ministry of Research (decisions #00762.02, #7822, #6463, #6466 and #19469, respectively), in compliance with French and European regulations.

### Cell lines

Vero cells (ATCC CRL-1586) were cultured at 37°C in Dulbecco’s Modified Eagle Medium (DMEM, Gibco) supplemented with 10% fetal bovine serum (FBS, Eurobio). C6/36 cells (ATCC CRL-1660) were cultured at 28°C in Leibovitz Medium (L-15 Medium, Gibco) supplemented with 10% FBS, 1% Non-Essential Amino Acids (Life Technologies) and 1% Tryptose Phosphate Broth (Life Technologies).

### Viruses

The FG15 Asian Zika virus (ZIKV) strain, isolated from a patient during ZIKV outbreak in French Guiana in December 2015, was obtained from the Virology Laboratory of the Institut Pasteur of French Guiana. The HD78788 African ZIKV strain, isolated from a human case in Senegal in 1991, was obtained from the Institut Pasteur collection. The KDH0026A DENV serotype 1 (DENV-1) strain, isolated from a patient in Thailand in 2010, was previously described (83). Viral stocks were prepared from supernatant of infected C6/36 cells, clarified by centrifugation at 800g and titrated on Vero cells by focus-forming assay (FFA). Stocks were kept at -80°C. The West Nile virus (WNV) strain IS-98-ST1 (or Stork/98) was obtained, cultured and used as described in Mashimo *et al.* (60). The Rift Valley Fever virus (RVFV) strain ZH548 was obtained, cultured and used as described in Tokuda *et al.* (84).

### Mouse experiments

All infection experiments were performed in a biosafety level 3 animal facility. Mice were maintained in isolators.

#### ZIKV and DENV systemic infection

CC mice received 2 mg of IFNAR-blocking mouse mAb (MAR1-5A3, BioXCell) by intraperitoneal (IP) injection one day before ZIKV or DENV infection (85). Groups of 6-8 week-old mice were inoculated IP with 10^7^ focus-forming units (FFUs) of ZIKV FG15 or 10^3^ FFUs of ZIKV HD78788, in 200µL PBS. For DENV infection, mice were anesthetized by IP injection with a solution of Xylazine (5 mg/kg) and Ketamine (80 mg/kg) and afterwards inoculated by intravenous (IV) injection in the retro-orbital sinus with 2.10^6^ FFUs of DENV-1 KDH0026A, in 100µL PBS. Survival and clinical signs were monitored daily for up to 14 days. Clinical signs were scored as follows: 0, no symptom; 1, ruffled fur; 2, emaciation, hunched posture and/or hypo activity; 3, hind limb weakness, prostration and/or closed eyes; 4, moribund or dead. Blood samples were collected at several time points from the retromandibular vein for plasma viral load assessment.

#### ZIKV intracerebral infection

Mice were anesthetized by IP injection with a solution of Xylazine (5 mg/kg), Ketamine (75 mg/kg) and Buprenorphine (0.03 mg/kg). Groups of 5-6 week-old mice were then inoculated by intracerebral (IC) injection in the right brain hemisphere with a 26-gauge needle affixed to a Hamilton syringe sheathed by a wire guard allowing no more than a 4-mm penetrance into the skull cavity, as described in (86). Mice received either 10^5^ FFUs of ZIKV FG15 in PBS or PBS alone, in a volume of 10µL. Survival and clinical signs were monitored daily for 6 days and euthanized for brain collection. Another cohort of mice (n=7-8 per strain) was infected similarly and monitored daily for 21 days to assess symptoms and survival.

#### WNV and RVFV infection

Groups of 8-12 week-old mice were inoculated IP with 10^3^ FFUs of WNV strain IS-98-ST1 or 10^2^ PFUs of RVFV strain ZH548. Survival and clinical signs were monitored daily for up to 14 days (RVFV) or 21 days (WNV).

### Mouse embryonic fibroblasts (MEFs) isolation and infection

MEFs were isolated from fetuses at day 13.5-14.5 of gestation, and cultured in DMEM supplemented with 10% FBS (Eurobio) and 1% penicillin/streptomycin (Gibco) at 37°C. MEFs were used until passage 2.

MEFs were plated at identical densities in culture dishes 24 hours before infection. MEFs were infected with ZIKV FG15 strain at a MOI of 5. After 2 hours of incubation at 37°C, the inoculum was replaced with fresh medium. Supernatants were collected at 24, 48 and 72 hours post-infection. Titration was performed by FFA in Vero cells.

### Mouse peritoneal macrophages (PMs) isolation and infection

Ten week-old mice were killed and PMs were collected by peritoneal lavage with 5 mL PBS (Gibco). The cell suspension was filtered on a 100 µm cell strainer and centrifuged at 800g for 10 minutes at 4°C. The cell pellet was re-suspended in serum-free RPMI 1640 medium (Gibco) and cells were plated in 96-well plates at desired density. After 1 hour incubation at 37°C, non-adherent cells were removed by 2 washes with PBS and fresh RPMI 1640 medium supplemented with 10% FBS and 1% penicillin/streptomycin was added to adherent cells.

Twenty-four hours after seeding, PMs were infected with ZIKV FG15 strain at a MOI of 5. After 2 hours of incubation at 37°C, the inoculum was replaced with fresh medium. Supernatants were collected at 24, 48 and 72 hours post-infection. Titration was performed by FFA in Vero cells.

### Mouse primary neurons isolation and infection

Primary neurons were prepared from mouse fetuses at day 16.5 of gestation. Isolated cortices were rinsed in HBSS medium (Gibco) and digested with 1 mg/mL Trypsin-EDTA (Gibco) and 0.5 mg/mL DNase I (Merck) in HBSS medium for 15 minutes at 37°C. B-27 supplement (Life Technologies) was added to inactivate Trypsin and mechanical dissociation of the cortices was performed by passages through a narrowed glass pipet. The cell suspension was centrifuged for 10 minutes at 200g and cell pellet was re-suspended in Neurobasal medium (Gibco) supplemented with 2% B-27, 0.2% L-glutamine (Gibco) and 1% penicillin-streptomycin-fungizone (Life Technologies). Cells were plated at identical densities in culture plates pre-coated with polyD-lysine (Merck) and Laminin (Merck).

Primary cultured neurons were infected with ZIKV FG15 strain at a MOI of 5 at 12 days of *in vitro* culture, for network maturation. After 2 hours of incubation at 37°C, the inoculum was replaced with fresh medium. Supernatants were collected at 24, 48 and 72 hours post-infection. Titration was performed by FFA in Vero cells.

### Focus-forming assay

Vero cells were seeded at 3.10^4^ per well in 100 µl complete medium (DMEM, FBS 10%) in 96-well plates. After overnight incubation at 37°C, medium was replaced with 40 µL of serial 10-fold dilutions of the samples, and 115 µL of methylcellulose overlay was added 2 hours later. After 40 hours incubation, culture medium was removed and cells were fixed with 100 µL/well of 4% paraformaldehyde for 20 minutes and permeabilized with a solution of 0.3% Triton and 5% FBS in PBS for 20 minutes. Cells were washed, incubated with a mouse mAb directed against ZIKV envelop protein (4G2, purified from the ATCC hybridoma) for 1 hour at 37°C (1/250^e^ in blocking buffer). Cells were further washed, incubated with secondary antibody (AlexaFluor-488-conjugated anti-mouse IgG, Invitrogen) for 45 minutes at 37°C and washed. Infected cell foci were counted using an ImmunoSpot CTL analyzer and viral titers were calculated from the average number of foci.

### Viral genome quantification by RT-qPCR

Blood samples were centrifuged to recover plasma from which viral RNA was extracted with the QIAamp Viral RNA Mini Kit (Qiagen). Brain samples were homogenized at 4°C in 1 mL of TRIzol reagent (Life Technologies) using ceramic beads and automated homogenizer (PreCellys). Total RNA was extracted according to manufacturer’s instructions. cDNA synthesis was performed using MMLV reverse transcriptase (Life Technologies) in a Bio-Rad *Mycycler* thermocycler. ZIKV and DENV cDNA were quantified by TaqMan quantitative PCR (qPCR) in a ViiA7 Instrument (Life Technologies) using standard cycling conditions. Primer sets adapted from previous works (87–89) were used to detect ZIKV and DENV RNA. ZIKV FG15 : forward, 5’-CCG CTG CCC AAC ACA AG-3’; reverse, 5’-CCA CTA ACG TTC TTT TGC AGA CAT-3’; probe, 5’-6FAM-AGC CTA CCT TGA CAA GCA ATC AGA CAC TCA A-MGB-3’ (Life Technologies). ZIKV HD78788: forward, 5’-AAA TAC ACA TAC CAA AAC AAA GTG GT-3’; reverse, 5’ -TCC ACT CCC TCT CTG GTC TTG-3’; probe, 5’-6FAM-CTC AGA CCA GCT GAA G-MGB-3’ (Life Technologies). DENV-1 KDH0026A: forward, 5’ –GGA AGG AGA AGG ACT CCA CA-3’; reverse, 5’-ATC CTT GTA TCC CAT CCG GCT-3’; probe, 5’ -6FAM CTC AGA GAC ATA TCA AAG ATT CCA GGG-MGB-3’ (Life Technologies). Viral load is expressed on a Log_10_ scale as viral genome copies per milliliter (plasma samples) or per total RNA microgram (brain samples) after comparison with a standard curve produced using serial 10-fold dilutions of a plasmid containing the corresponding fragment of ZIKV genome.

### Western Blot analysis

MEFs (5.10^6^) were pre-incubated with IFNAR1-blocking antibody (MAR1-5A3, BioXCell) for 7 hours and then stimulated, or not, with 300 IU/mL mouse IFN-α (Miltenyi Biotec) for 15 minutes. MEFs were detached and centrifuged at 300xg for 5 minutes and cell pellet was re-suspended in cold PBS. MEFs were then lyzed into extraction buffer (10 mM Tris–HCl, pH 7.5, 5 mM EDTA, 150 mM NaCl, 1% NP40, 10% glycerol, 30 mM NaP, 50 mM NaFluoride) completed with protease inhibitor (Complete, EDTA free, Roche) and phosphatase inhibitors (phosStop easy pack, Roche) with 2.5 UI of benzonase nuclease (Sigma). Lysates were incubated on ice for 30 minutes and non-soluble fraction was separated by centrifugation. Protein concentrations were determined by Bradford assay, and equal amounts of protein were further used. Protein denaturation was performed in Laemmli buffer at 95°C for 5 minutes. After separation on a 12% polyacrylamide gel (Biorad), proteins were transferred on Immun-Blot polyvinylidene difluoride (PVDF) membrane (Biorad) and incubated overnight with the following antibodies: Anti-Phospho-Stat1 Tyr701 (1/1,000^e^, #9167, Cell Signaling), Anti-Total Stat1 N-terminus (1/500^e^, # 610115, BD Biosciences), Anti-αTubulin (1/8,000^e^, # T5168, Merck). Membranes were incubated for 1.5 hour at room temperature with an anti-mouse or an anti-rabbit IgG horseradish peroxidase-linked secondary antibody (1/10,000^e^, NA931 and NA934V, Amersham) and signals were visualized using autoradiography.

### Histopathology

After necropsy, brain was removed, fixed for 48-72 hours in 10% neutral-buffered formalin and embedded in paraffin; 4µm-thick sections were stained in hematoxylin-eosin. Morphology of microglial cells was assessed by immunohistochemistry using rabbit anti-Iba1 primary antibody (# 01919741, Wako chemical, dilution 1:50) as previously described (90). Sections were analyzed by a trained veterinary pathologist in a blind study on coded slides.

### Genetic analysis

Broad sense heritability was calculated as previously described (59).

Plasma viral load at days 2 and 6 p.i. and plasma viral load decrease, measured on 159 mice from 35 CC strains (average of 4.5 mice per strain), were used in quantitative trait locus (QTL) mapping using the rqtl2 R package (91) and GigaMUGA genotypes of CC founders and CC strains available from http://csbio.unc.edu/CCstatus/CCGenomes/#genotypes. Genome scan was performed using the scan1 function with a linear mixed model using a kinship matrix. Statistical significance levels were calculated from 1,000 permutations.

Genotype-phenotype associations for specific genes (*Ifnar1*, *Oas1b*) were tested by Kruskal-Wallis test using the founder haplotype as the genotype.

### Statistical analysis

Statistical analyses were performed using R software (v3.5.2). Kaplan-Meier survival curves were compared by logrank test. Two-way ANOVA was used for testing mouse strain and sex effects on plasma viral load at day 2 p.i. (FIG 2E). Student’s *t*-test was used to compare viral loads in tissues, except when data showed heterogeneous variance between groups, in which case we used Kruskal-Wallis and Wilcoxon non-parametric tests. These tests were also used for assessing mouse strain effect on plasma viral load and on plasma viral load decrease (FIG 3). Pearson’s coefficient was used for the correlation between plasma viral load at days 2 and 6 p.i. (FIG 3B) and for the correlation between measurements of plasma viral load by FFA and RT-qPCR (FIG 2D). Student’s *t*-test was used to compare viral titers between strains in *in vitro* experiments. P-values < 0.05 were considered statistically significant.

## NOTES

We are grateful to the Virology Laboratory of the Institut Pasteur of French Guyana (National Reference Center for Arboviruses) for providing the FG15 ZIKV strain and Valérie Choumet for providing the IS-98-ST1 WNV strain. We thank Thérèse Couderc and Claude Ruffié for providing B6-*Ifnar1* and 129-*Ifnar1* mice, Laurine Conquet, Laetitia Joullié and Marion Doladilhe for technical help, Magali Tichit for histopathology techniques, Isabelle Lanctin and Jerôme Le Boydre for careful breeding of CC mice, and the animal facility staff for animal care in biocontainment units (DTPS-C2RA-Central Animal Facility platform). We are grateful to Jean Jaubert, Michel Cohen-Tannoudji and Aurore Vidy-Roche for useful discussions throughout the project, and to Rachel Maede for editorial suggestions.

The authors declare no competing interests.

This work was supported by a grant from the French Government’s Investissement d’Avenir program, Laboratoire d’Excellence “Integrative Biology of Emerging Infectious Diseases” (grant n°ANR-10-LABX-62-IBEID). C.M. was supported by a fellowship from the grant n°ANR-10-LABX-62-IBEID.

